# A near complete genome assembly of chia assists in identification of key fatty acid desaturases in developing seeds

**DOI:** 10.1101/2022.08.15.504044

**Authors:** Leiting Li, Jingjing Song, Meiling Zhang, Shahid Iqbal, Yuanyuan Li, Heng Zhang, Hui Zhang

## Abstract

Chia is an annual crop whose seeds have the highest content of α-linolenic acid (ALA) of any plant species. We generated a high-quality assembly of the chia genome using circular consensus sequencing of PacBio. The assembled six chromosomes are composed of 21 contigs and have a total length of 361.7 Mb. Genome annotation revealed a 53.5% repeat content and 35,850 protein-coding genes. Chia shared a common ancestor with *Salvia splendens* ~6.1 million years ago. Utilizing the reference genome and two transcriptome datasets, we identified candidate fatty acid desaturases responsible for ALA biosynthesis during chia seed development. Because the seed of *S. splendens* contains significantly lower proportion of ALA but similar total contents of unsaturated fatty acids, we suggest that strong expression of two *ShFAD3* genes are critical for the high ALA content of chia seeds. This genome assembly will serve as a valuable resource for breeding, comparative genomics, and functional genomics studies of chia.

## Introduction

Chia (*Salvia hispanica* L.) is an annual herbaceous crop belonging to the family of Lamiaceae, also commonly known as the mint family. Chia is native to central America and is believed to serve as a staple crop of the Aztec in pre-Columbian times (Valdivia-López and Tecante, 2015). Chia is currently cultivated for its seeds in Central and South America. Chia produces oily seeds with an oval shape and a diameter of ~2 mm. Thanks to its superior nutrient compositions, the chia seed is a trending functional food ingredient (Muñoz et al., 2013; Cassiday, 2017). Chia seeds contain 30-40% total lipids, of which α-linolenic acid (ALA; C18:3, n-3), linoleic acid (LA; C18:2, n-6), and oleic acid (C18:1, n-9) account for ~60%, ~20%, and ~10% respectively (Ciftci et al., 2012; Kulczynski et al., 2019). ALA is an essential fatty acid (i.e., cannot be synthesized by human body) and up to 8-21% and 1-9% of ALA intake can be respectively converted to eicosapentaenoic acid (EPA; C20:5, n-3) and docosahexaenoic acid (DHA; C22:6, n-3) in the human body (Baker et al., 2016; Shahidi and Ambigaipalan, 2018). Studies indicate that these n-3 fatty acids are important for human development and growth (Li et al., 2019). The recommended Adequate Intake (AI) of ALA is 1.6 g/day for men and 1.1 g/day for women (Burns-Whitmore et al., 2019). In addition, a low n-6:n-3 ratio, as in the case of chia seeds, in the diet helps reduce inflammation (Simopoulos, 2002, 2002; Lands, 2014). Chia seeds also have high contents of dietary fiber (up to 34.4%), proteins (16.5-24.2%), vitamin B3, multiple minerals (such as calcium, phosphorus, potassium, and ion), and antioxidants (Kulczynski et al., 2019). Because of these properties, chia seeds are increasingly used as an ingredient in food industry and restaurants.

In plants, fatty acid (FA) biosynthesis takes place within the plastid, where acetyl-coenzyme A (acetyl-CoA) is used as the main carbon donor for the initiation and elongation of acyl chains (Ohlrogge and Browse, 1995; Li-Beisson et al., 2013). During the elongation, fatty acids remain covalently attached to acyl carrier proteins (ACPs), which serve as a cofactor for FA biosynthesis. The fatty acids biosynthesis cycle is usually terminated when the acyl chain reaches 16 or 18 carbons in length, and two principal types of acyl-ACP thioesterases, FatA and FatB, hydrolyze acyl-ACP and release the corresponding FAs. Desaturation of common fatty acids (C16 and C18) begins at the C-9 position (Δ9) and progresses in the direction of the methyl carbon of the acyl chain. Thus, the conversion of stearic acid (C18:0) to α-linoleic acid (C18:3^Δ9,12,15^) involves the sequential action of three desaturases, including the stearoyl-ACP desaturase, the oleate desaturase, and the linoleate desaturase. In the model plant Arabidopsis, genetic analyses have identified the main enzymes with specific FA desaturase activities. While all the other FA desaturases are membrane-bound enzymes, the family of acyl-ACP desaturases (AADs) are stromal soluble enzymes that use stearoyl-ACP (C18:0) or palmitoyl-ACP (C16:0) as the substrate. The Arabidopsis genome encodes 7 AADs (Kachroo et al., 2007), named as FAB2 (FATTY ACID BIOSYNTHESIS 2) and AAD1-6. Genetic analyses indicate that FAB2, AAD1, ADD5, and AAD6 are redundant Δ9 stearoyl-ACP desaturases (SADs) (Kazaz et al., 2020), while AAD2 and AAD3 function as Δ9 palmitoyl-ACP desaturases (PADs) (Troncoso-Ponce et al., 2016). Further desaturation of oleic acids (C18:1^Δ9^) may take place within the plastid or the endoplasmic reticulum (ER). In the plastid, the oleic acids are incorporated into multiple types of glycerophospholipids and converted to C18:3 by FAD6 (FATTY ACID DESATURASE 6) and FAD7/8. Alternatively, the oleic acid may be exported and enters the acyl-CoA pool in the cytosol. The C18:1-CoA can be imported into ER, where it is incorporated into phosphatidylcholine (PC) and becomes sequentially desaturated by FAD2 and FAD3, which respectively prefer PC with C18:1 and C18:2 as the substrate. During seed development, the desaturated PCs are further converted to diacylglycerol (DAG) and triacylglycerol (TAG), the latter of which is the main form of storage lipids in the oil body of seeds.

In this study, we assembled a high-quality chia genome using accurate consensus long reads (PacBio HiFi reads). The six chia chromosomes are composed of 21 main contigs, with telomere repeats at 8 ends of the chromosomes. Utilizing this highly accurate and complete genome and a published seed development transcriptome, we identified the main ER-localized linoleate desaturases that underlie the extremely high ALA content in chia seeds.

## Results

### Genome assembly

We selected a chia cultivar with a Mexico origin (**Supplemental Figure 1**) for the assembly of the genome. About 24.7 Gb of circular consensus sequencing reads with an average read length of 16.1 kbp were generated from a single sequencing cell (**Supplemental Figure 2**). K-mer-based analyses of the HiFi reads estimated the nuclear genome to be ~352.7 Mb in size (**Supplemental Figure 3**).

We performed genome assembly using the hifiasm assembler (Cheng et al., 2021). The initial assembly was 388.0 Mb, consisting of 666 contigs with a N50 length of 21.8 Mb and an L50 number of 7, indicating a high contiguity of the assembly. The longest 21 contigs have a total length of 361.7 Mb and a minimum length of 1.7 Mb, while other contigs are significantly shorter, 636 of which have lengths shorter than 150 kbp (**Figure 1A**). The average HiFi read depth on the 21 long contigs varies between 43 and 58, which are around the 54-fold coverage of the nuclear genome calculated from the k-mer distribution (**Figure 1A**; **Supplemental Figure 3**). In contrast, the rest 645 contigs have a read coverage varying from 0 to 557, suggesting that they originate either from fragments of highly repetitive regions or from the high-copy organellar genomes.

**Figure 1.**
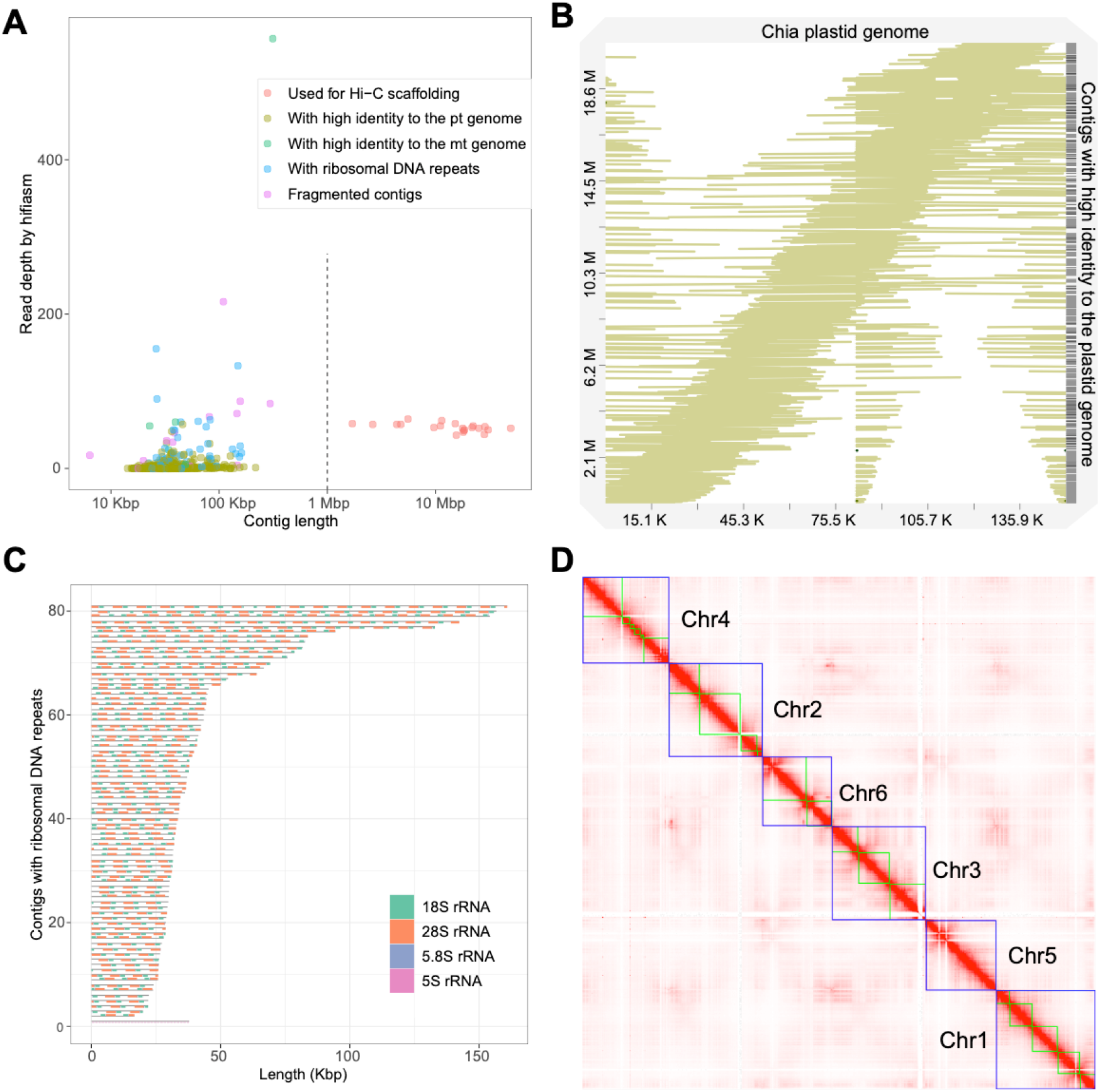
Assembly of the chia genome. **A)** Dotplot showing the contig length and the read depth of the initial assembly. Contigs were classified into five categories based on the length, the read depth and their origins, as indicated in the legend. **B)** Alignment of 538 initial contigs onto the chia plastid genome. **C)** Structure of the 81 contigs containing ribosomal RNA repeats. **D)** Hi-C contact map of the chia nuclear genome. Blue boxes indicate grouped pseudochromosomes, whereas green boxes indicate contigs.

We next analyzed the plastid and mitochondrion genomes. From the initial assembly, we identified a circular contig (ptg000033c) that has a length of 313,444 bp and an average read coverage of 557 folds. Genome annotation identified 151 mitochondrion-encoded genes, including 21 transfer RNAs, 6 ribosomal RNAs (rRNAs), and 124 protein-coding genes (**Supplemental Figure 4**), indicating that this contig is the complete mitochondrion genome. We also identified 4 other contigs that show high sequence identity (100%) to the mitochondrion genome (**Supplemental Figure 5**). We reason that these contigs may represent mitochondrial genome fragments recently transferred to the nuclear genome.

We could not identify a single contig representing the plastid genome from the initial assembly. We thus assembled the plastid genome using Illumina short reads and the GetOrganelle software (Jin et al., 2020). The plastid genome has a length of 150,956 bp and 132 genes, including 87 protein-coding genes, 37 tRNA genes, and 8 rRNA genes (**Supplemental Figure 6**). Surprisingly, we found that 538 out of the 666 initial contigs could be mapped to the plastid genome with high coverage (>99%) and high identity rate (>99%) (**Figure 1B**). These contigs are short in length (14.2 to 217.6 kb) and most of them have low HiFi read coverage (with 530 contigs below 19-fold coverage) (**Figure 1A**). These plastid-originated contigs likely represent misassembled plastid genome fragments and/or nuclear genome fragments with a plastid origin. The total length of these contigs was 20.7 Mb, accounting for most of the excessive part of the assembly compared to the predicted genome size.

Excluding the organellar-originated 543 contigs and the 21 high-confidence nuclear contigs, the rest 102 contigs have a total length of 5.2 Mb. Ribosomal RNA (rRNA) repeats were identified in 81 of these contigs, indicating they were originated from genomic regions with high copy number of rRNA genes. Except for one contig mainly composed of 73 repeats of 5S rRNA, other contigs had a basic repeat unit of a “18S-5.8S-28S” structure with the copy number varied from 2 to 17 (**Figure 1C**). Considering the nuclear origin of most sequences, the 102 contigs were concatenated as Chr0.

We next used the 21 high-confidence nuclear contigs for Hi-C scaffolding. Based on ~180x (63.8 Gb) of Hi-C sequencing data, we clustered and ordered the 21 contigs into six pseudochromosomes, whose sizes ranged from 47.8 Mb to 69.1 Mb (**Figure 1D; Figure 2; Table 1**). Chr5 was composed of a single contig while Chr4 contained the largest number (6) of contigs. The total length of the six pseudochromosomes was 361.7 Mb. The final v1 assembly (Shi_PSC_v1) of the chia genome composed of 9 sequences, seven of which (Chr0-Chr6) represent the nuclear genome, one for the mitochondrion genome, and one for the plastid genome.

**Figure 2.**
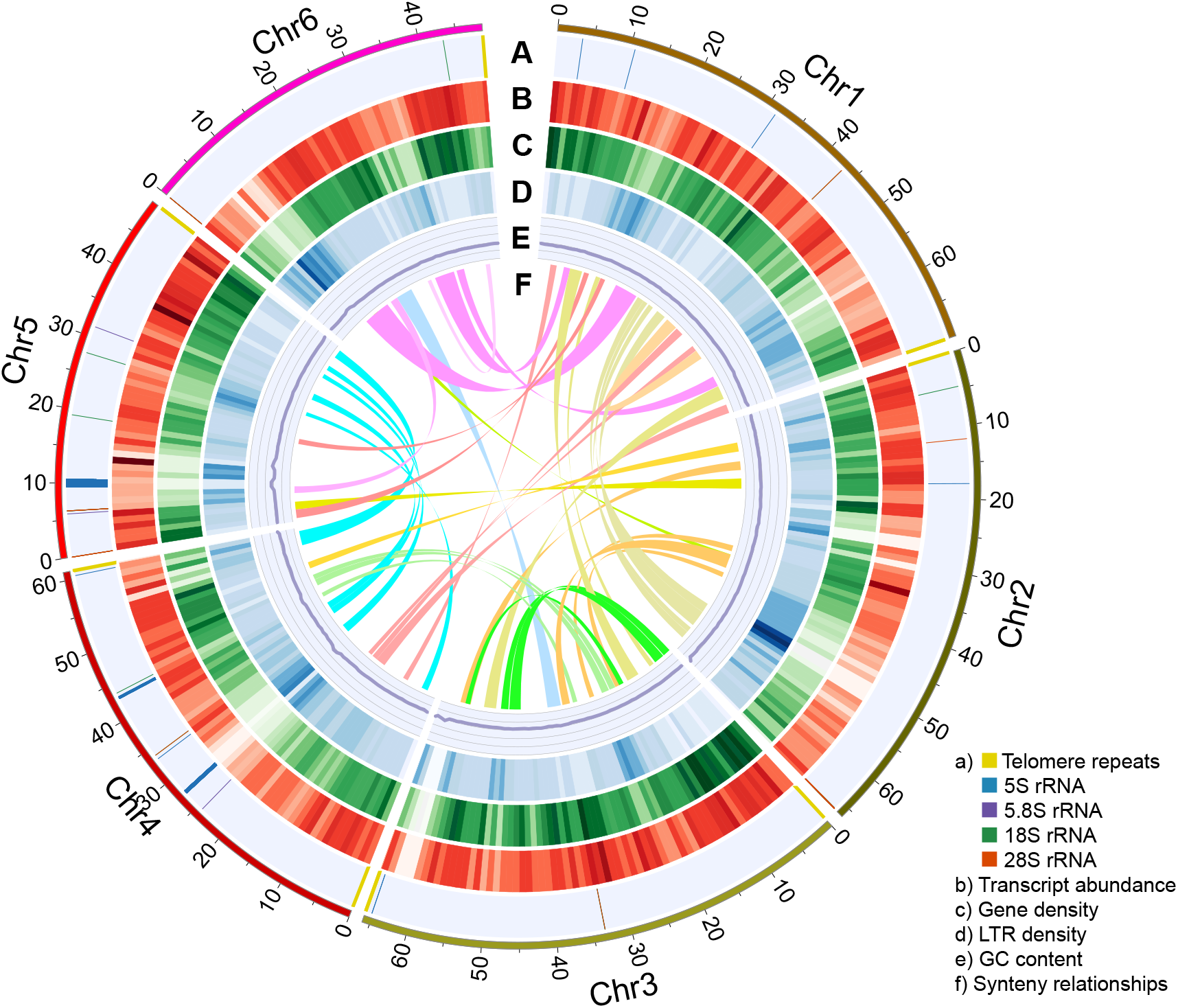
The nuclear genome of Shi_PSC_v1. Each ring indicates specific features of the nuclear genome. Data from non-overlapping 1-Mb windows were graphed: **A)** Position of telomere repeats and ribosomal RNA genes; **B)** Average transcript abundance; **C)** Gene density; **D)** LTR density; **E)** GC content; **F)** Synteny blocks >1 Mb in length.

**Table 1.**
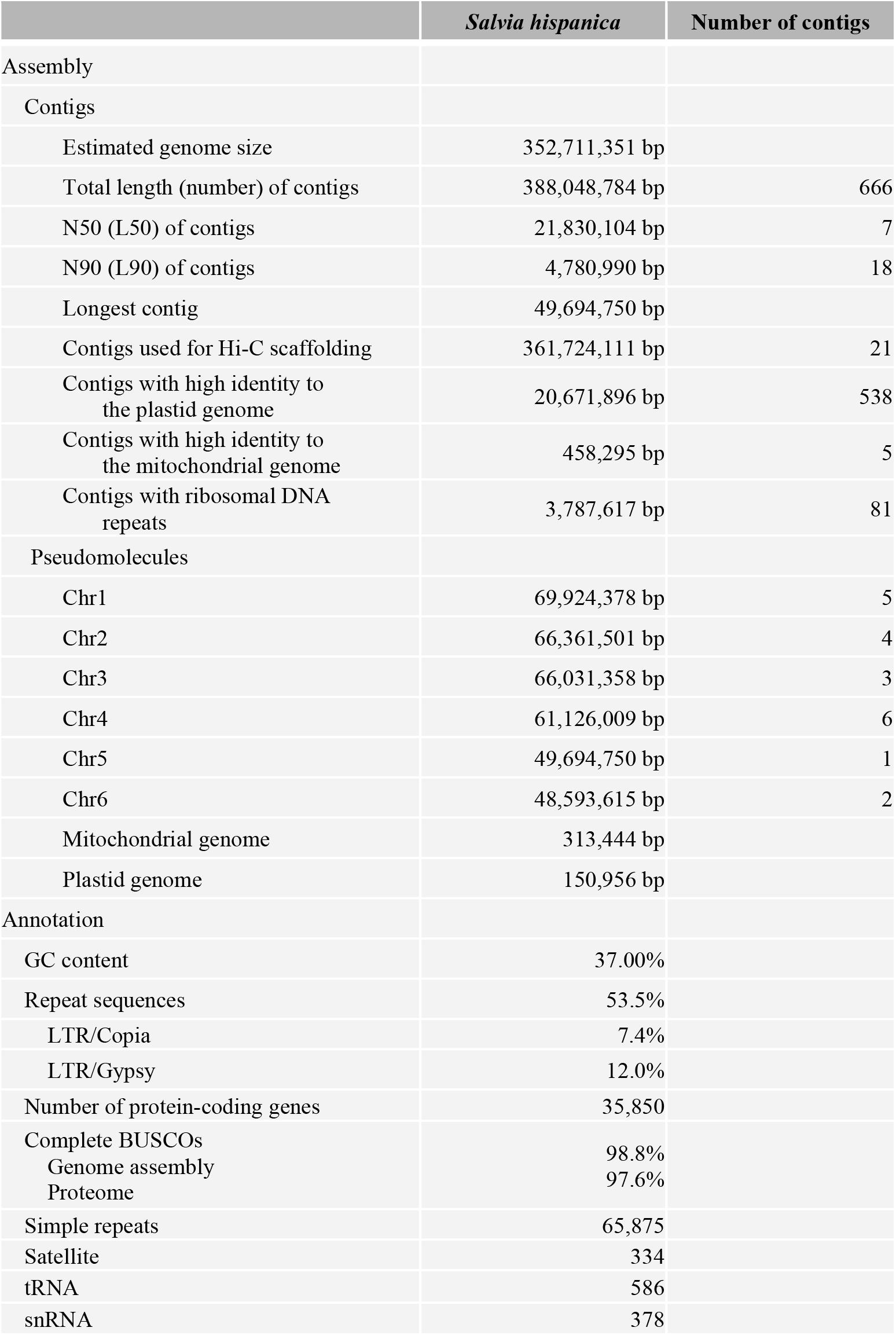
Summary of chia genome assembly and annotation

### Evaluation of genome assembly

We next evaluated the quality of the genome assembly using LTR Assembly Index (LAI) (Ou et al., 2018), Benchmarking Universal Single-Copy Orthologs (BUSCO) (Manni et al., 2021), Merqury (Rhie et al., 2020) and Illumina short reads. The whole genome had an LAI of 15.78, which was around the same level as the TAIR10 assembly of *Arabidopsis thaliana*, and could be considered as the reference level (Ou et al., 2018). The complete BUSCO of the chia genome assembly was 98.8%, indicating a high completeness of the gene space. Merqury compares k-mers from the assembly to those found in unassembled HiFi reads to estimate the completeness and accuracy. The completeness and quality value (QV) of Shi_PSC_v1 were 97.3 (out of 100) and 66.5 (>99.99% accuracy) respectively. Mapping of the Illumina short reads (**Supplemental Table 1**) against the chia genome assembly also revealed very high read mapping rate (99.9%) and a low apparent error rate (0.27%).

### Genome annotation

For genome annotation, we first identified repetitive sequences in the Shi_PSC_v1 assembly. The analysis revealed that chia nuclear genome had a repeat content of 53.5% (**Table 1**). Similar to most plant genomes, retrotransposons accounts for the majority of the repetitive sequences of the genome. About half of the repeats were characterized as long terminal repeats (LTRs), with Gypsy and Copia being the main types. Besides, 65,851 simple repeats, 334 satellite sequences, 573 transfer RNAs (tRNAs) and 378 small nuclear RNAs (snRNAs) were also identified in the chia genome (**Supplemental Table 2**).

The repeat-masked assembly was then used for gene model prediction. Based on evidence from *ab initio* prediction, expressed sequence tags (ESTs), and homologous protein sequences, a total of 35,850 protein-coding genes were annotated. Additionally, we also examined whether telomere signals were present at the end of each pseudochromosome. The results showed that all the six pseudochromosomes contain telomere repeats. Telomere repeats were detected at both ends of Chr3 and Chr4, and one end of Chr1, Chr2, Chr5, and Chr6 (**Figure 2A**).

The complete BUSCO score of the protein sequences was 97.6%, close to the BUSCO score of the genome assembly (98.8%). Functional annotation showed that Gene Ontology (GO) terms (Gene Ontology, 2021), Pfam domains (Mistry et al., 2021), and InterPro families (Blum et al., 2021) were assigned to 58.9% (21,125), 72.0% (25,799), and 79.2% (28,405) of the protein-coding genes. In total, AHRD (Automated assignment of Human Readable Descriptions) function names were assigned to 89.5% (32,089) of the protein-coding genes (Boecker, 2021) (**Supplemental Table 3**). These metrics indicate high quality of the genome annotation.

### Evolution of the chia genome

To understand the evolution of the chia genome, we selected five other species from the family of Lamiaceae, including three from the genus of *Salvia*, together with three species of Asterids and *Arabidopsis thaliana* for the orthology analysis (**Figure 3A**). A species tree constructed using orthologs shared in all analyzed species by STAG (Emms and Kelly, 2018) confirmed a close relationship between chia and *S. splendens*, as well as *S. bowleyana* and *S. miltiorrhiza* (**Figure 3A**). After calibrating divergence time using data retrieved from the TimeTree database (Hedges et al., 2015), chia was estimated to diverge with *S. splendens* ~6.2 million years ago (MYA) and the four *Salvia* species have a common ancestor ~21.8 MYA. The protein-coding genes of chia were assigned to 17,158 families. Relative to the common ancestor of chia and *S. splendens*, expansion in 528 families and reduction in 2,344 families were observed in chia (**Figure 3A**). In contrast, *S. splendens* had 8,777 expanded families and a large number of 2-copy gene families (**Figure 3B**). This is consistent with its recent tetraploidization event (Jia et al., 2021). Among the ten species analyzed, 8,812 families were shared while between 265 and 1,147 families were unique for each species (**Figure 3C**). Among the 720 gene families (2,529 genes) unique to chia, 72.6% of them were comprised of 2 or 3 members (**Supplemental Figure 7**) and the largest one contained 36 members. GO enrichment analysis was performed for genes in these chia-specific gene families. The results showed that the top enriched GO term in the category of biological process was “defense response” (GO:0006952) (**Supplemental Figure 8**), suggesting their potential roles in the environmental adaptation of chia. In addition, “acyl-[acyl-carrier-protein] desaturase activity” (GO:0045300) in the category of molecular function was enriched (**Supplemental Figure 9**). This expanded family mainly includes orthologs genes of *AtFAB2* (**Supplemental Table 3; Supplemental Figure 10**), the stearoyl-ACP (C18:0) or palmitoyl-ACP (C16:0) desaturases of Arabidopsis.

**Figure 3.**
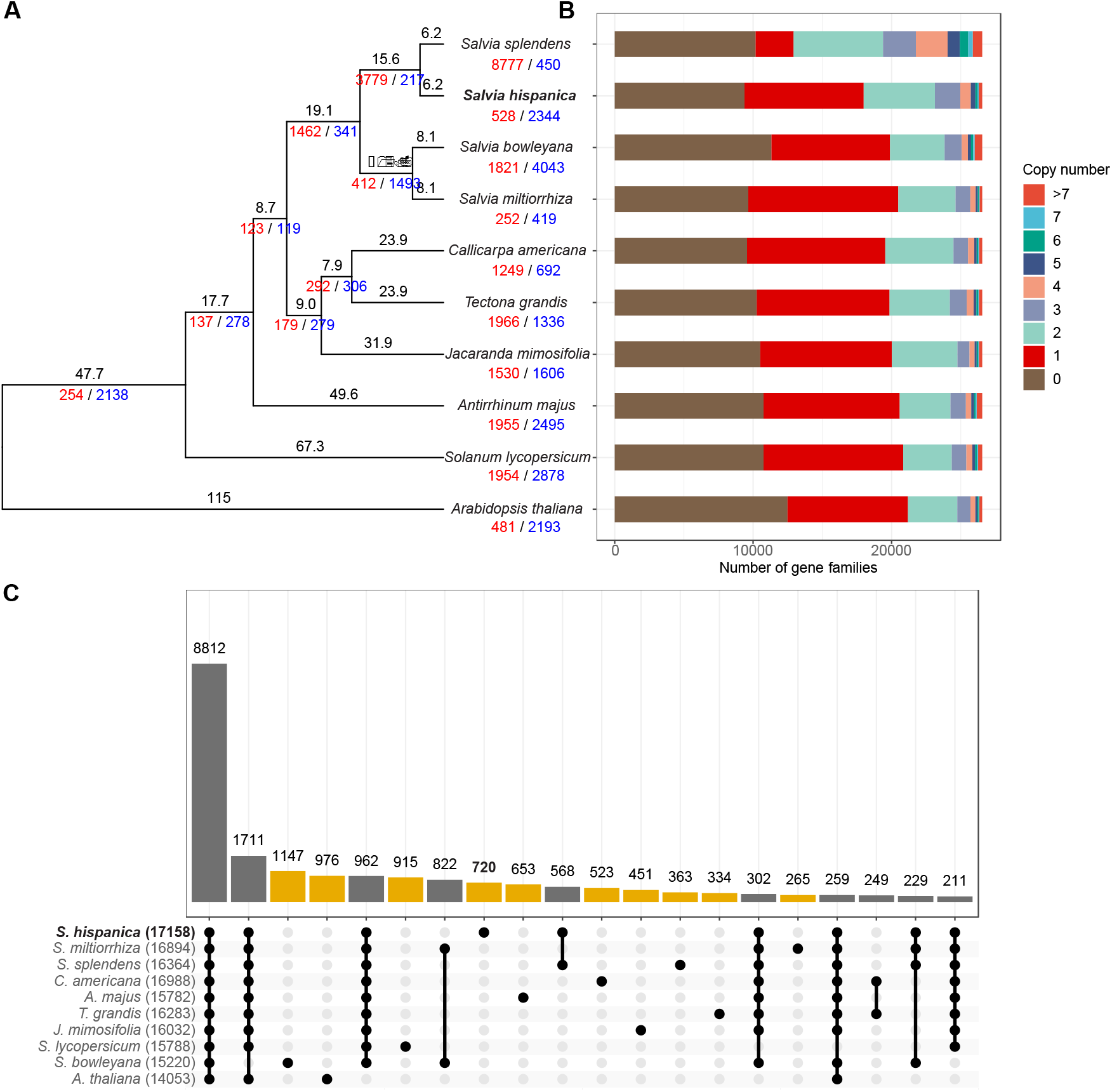
Evolution of the chia genome. **A)** Phylogenetic tree for chia and 9 other plant species. Numbers of expanded and contracted gene families were indicated by red and blue numbers at each branch point. Branch length indicate the estimated divergence time in million years ago. **B)** Numbers of gene families with different copy numbers in each plant species. **C)** Upset plot indicating the number of gene families shared by different species. Yellow bars indicate species-specific gene families.

To investigate the whole-genome duplication events of chia, we performed intra-genome synteny analysis. In total, 323 synteny blocks with an average of 20.5 homologous gene pairs per block were identified (**Figure 2F**). The distribution of synonymous substitution rates (Ks) of these gene pairs revealed a single Ks peak at ~0.26 (**Supplemental Figure 11**), which was consistent with the whole genome duplication (WGD) event prior to the tetraploidization event of *S. splendens* (Jia et al., 2021). This indicates that this WGD event occurred before the divergence of chia and *S. splendens*.

### Identification of genes involved in ALA biosynthesis

We next sought to identify genes underlying the high ALA content in chia seeds. We used kofamKOALA (Aramaki et al., 2020) to identify homologous genes of the lipid biosynthesis pathway (ko01004 of KEGG) in the chia genome (**Supplemental Figure 12; Supplemental Table 4**). We focused on genes encoding fatty acid desaturases. The analysis revealed 2 homologs of *AtFatA* (K10782), 6 homologs of *AtFatB* (K10781), 14 genes of the *AAD* family (K03921), 2 homologs of *AtFAD2* (K10256), and 4 homologs of *AtFAD3/7/8* (K10257), among others (**Supplemental Figure 12**). Further phylogenetic analyses separated *AtFAD3/7/8* (K10257) into 2 branches, each containing 2 orthologs of *AtFAD3*, and *AtFAD7/8* (**Figure 4A**; **Supplemental Figure 13**). Multiple sequence alignment (**Supplemental Figure 14**) indicated that *AtFAD7/8* and their orthologs in chia contain extra N-terminal sequences (plastid transit peptides) compared to the *AtFAD3* branch, consistent with their predicted localization in the plastid (Xue et al., 2018).

**Figure 4.**
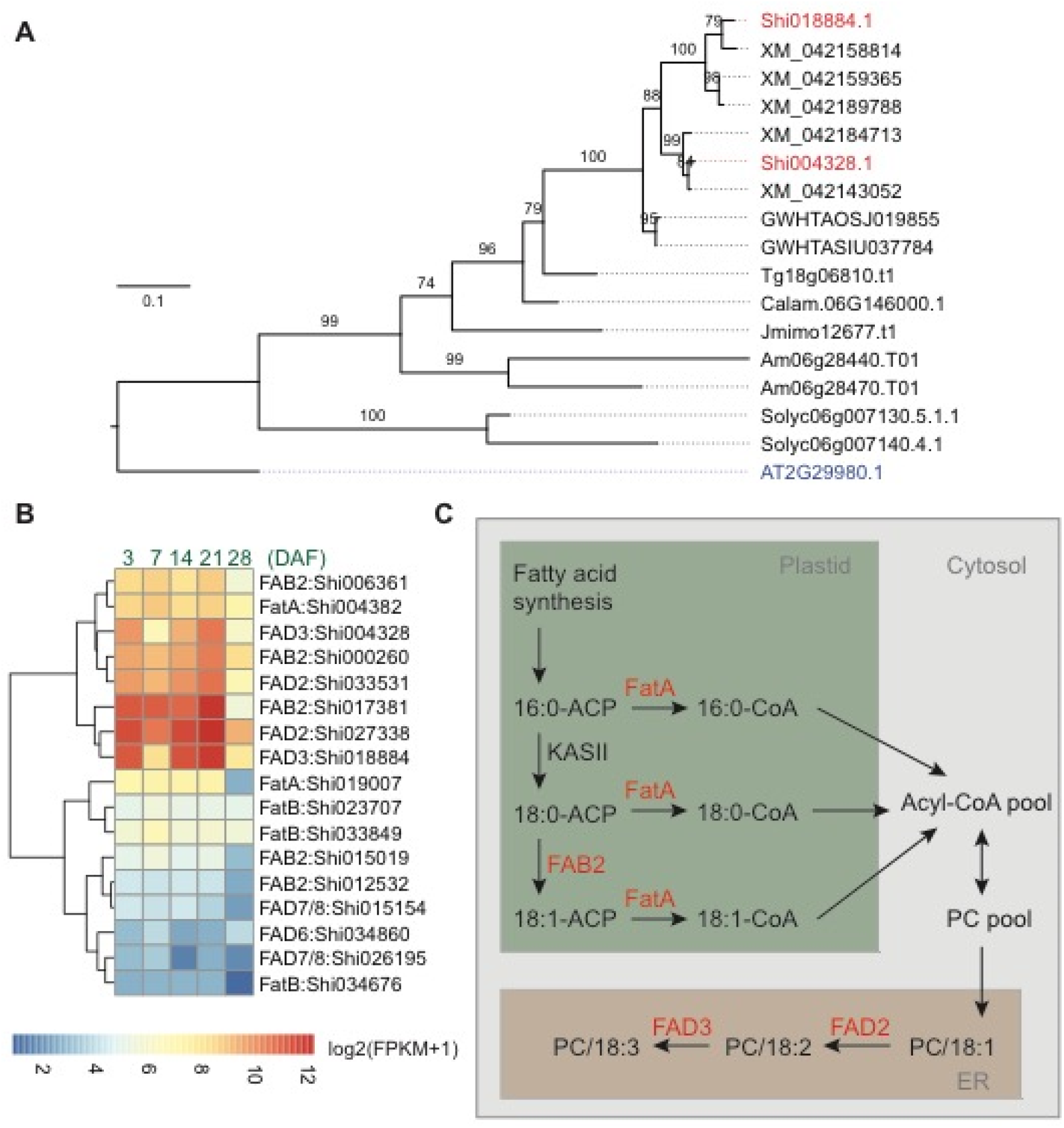
Identification of critical genes involved in fatty acid biosynthesis of chia seeds. **A**) Phylogenetic tree of the *FAD3* genes. Shi: *Salvia hispanica;* AT: *Arabidopsis thaliana;* XM: *Salvia splendens;* GWHTAOSJ: *Salvia miltiorrhiza;* GWHTASIU: *Salvia bowleyana;* Tg: *Tectona grandis*; Jmimo: *Jacaranda mimosifolia*; Calam: *Callicarpa americana*; Am: *Antirrhinum majus;* Solyc: *Solanum lycopersicum*.**B)** Expression pattern for *FatA, FatB, FAB2, FAD2, FAD3, FAD7/8*, and *FAD6* genes in developing chia seeds. DAF: Days after flower opening; FPKM: Fragments Per Kilobase of transcript per Million mapped reads. Only genes with maximum FPKM > 1 in seed samples were included in the plot. **C)** A model for the biosynthesis of ALA in the chia genome. PC: phosphatidylcholine; ER: endoplasmic reticulum.

We utilized two published transcriptome dataset to help identify candidate ALA biosynthesis genes in the chia genome, one covering 13 different tissues or developmental stages of chia (Gupta et al., 2021) and one covering five different time points of chia seed development (3, 7, 14, 21, and 28 days after flower opening (DAF)) (Sreedhar et al., 2015). We reason that the candidate genes should be expressed at significantly higher levels compared to their counterparts during seed development. Indeed, we found that *Shi004382* (*ShFatA*), *Shi017381*, *Shi000260*, and *Shi006361* (*AtFAB2* orthologs), *Shi027338* and *Shi033531* (*AtFAD2* orthologs), and *Shi018884* and *Shi004328* (*AtFAD3* orthologs) are highly expressed in developing chia seeds, and their expression levels are decreased in the 28 DAF sample (**Figure 4B**). These genes are also expressed at significantly higher levels in developing seeds compared to other chia tissues/organs (**Supplemental Figure 12**). Although FAB2 homologs have either SAD or PAD activity, studies in Arabidopsis indicate that a single amino acid change (Tyr to Phe) is sufficient to confer PAD activity to AtFAB2 (SAD) (Troncoso-Ponce et al., 2016). The residue is predicted to locate at the bottom part of the substrate channel and the bulkier lateral chain of Phe may reduce the substrate binding pocket to better accommodate C16-ACP substrates. Multiple sequence alignment indicated that the highly expressed FAB2 homologs (*Shi017381, Shi000260*, and *Shi006361*) in chia seeds have a Tyr residue at the corresponding position, suggesting that they function as SADs (**Supplemental Figure 15**). In contrast, the two orthologs (*Shi015154* and *Shi026195*) of *AtFAD7/8*, the plastid localized omega-3 desaturase of Arabidopsis, were expressed at low to medium levels (FPKM values between 1.7 – 18.6) in developing seeds (**Figure 4B**; **Supplemental Figure 12**). In fact, the most highly expressed genes in developing chia seeds also contain multiple FA biosynthesis-related genes, such as genes encoding acyl carrier proteins (*Shi029800*, *Shi029801* and *Shi008432*), oil body-associated proteins (*Shi002948* and *Shi002148*), and lipid-transfer proteins (*Shi014949* and *Shi010250*) (**Supplemental Table 5**). These results suggest a biosynthetic pathway involving plastid and ER localized enzymes, including *ShFAB2, ShFatA, ShFAD2* and *ShFAD3*, is responsible for the high ALA content in chia seeds (**Figure 4C**). Despite copy number variations were identified in some of these genes (**Supplemental Table 4**), we suggest that strong expression of fatty acid desaturase genes, particularly the ER localized FAD3s, are responsible for the high ALA content in chia seeds.

## Discussion

*De novo* assembly of plant genomes has been greatly facilitated by the advancement of third-generation sequencing technologies that produce single-molecule long reads without the need of polymerase chain reactions. Commercially available 3^rd^-generation sequencing platforms suffer from high error rate of the raw reads (usually between 10-15%). The circular consensus sequencing (CCS) mode of PacBio significantly reduced consensus error rate by sequencing the same DNA insert multiple times. With carefully selected sizes of the DNA insert, a balance of sequencing length and accuracy can be achieved. In the current study, we performed CCS sequencing of the chia genomic DNA with a single SMRT cell, which produces 24.7 Gb of CCS data with median quality value of 31. The initial assembly included 666 contigs, while our analyses indicated that 623 of them originated from the organellar genomes or ribosome RNA repeats (**Figure 1**). The top 21 contigs have a total length of 361.7 Mb, which is slightly larger than the estimated genome size of 352.7 Mb based on k-mer analysis. Consistent with this high completeness of the nuclear genome, telomere repeats were identified at one or both ends of each of the six pseudochromosomes and rRNA repeats were identified in multiple chromosomes (**Figure 2**). Collapsing of repetitive regions was a common problem for *de novo* assembly of genomes with high repeat contents using longer but non-CCS PacBio reads. We did not observe similar phenomenon during the assembly of the chia genome. We reason that improved accuracy of the CCS mode helps resolving highly complex regions of the genome unless the repeat unit exceeds the read length, or the repeat sequences are highly similar.

Through phylogenetic and gene expression analyses, we identified candidate genes underlying high ALA contents of chia seeds. Two copies each of *ShFAB2, ShFAD2*, and *ShFAD3* exhibit very similar expression patterns (**Figure 4B**), suggesting these enzymes act together to promote the ALA content in chia seeds. This is consistent with the reported substrate channeling between FAD2 and FAD3 (Lou et al., 2014). Mature chia seeds have a lipid content of ~35%, of which up to 64% are ALA, the highest among all plant species (Muñoz et al., 2013; Kulczynski et al., 2019). Compared to its close relative, *S. splendens*,whose seeds were reported to have a ALA content of 34.5% and a LA content of 31.3% (Joh et al., 1988), the total content of ALA and LA of chia seeds are similar, suggesting that the elevated conversion rate from LA to ALA is the main event that drives high ALA content in chia seeds. In support of the idea that FAD3 is a rate limiting step in ALA biosynthesis, it was shown that overexpression of the rice *FAD3* gene is sufficient to increase the ALA content in seeds by ~28 fold (Liu et al., 2012). In addition to chia, seeds of flax (*Linum usitatissimum*) and perilla (*Perilla frutescens*) also have a relative ALA content around 60% (Ciftci et al., 2012). Although the genetic bases underlying their high ALA content remains to be determined, convergent high ALA contents in these species indicate that increasing omega-3 contents in seeds involve limited number of steps during evolution. This suggests a promising future for improving lipid composition in grains through transgenic or genome editing approaches.

## Materials and methods

### Library preparation and sequencing

Chia seeds were surface sterilized and grown in ½ MS medium supplemented with 0.7% agarose in a Percival growth chamber. Genomic DNA was extracted from two-week-old seedlings for genome survey sequencing and accurate consensus long-read sequencing (HiFi sequencing). The genome survey library was prepared and sequenced at the Genomics Core Facility of Shanghai Center for Plant Stress Biology following standard protocols. A 15-kb PacBio HiFi sequencing library were constructed and sequenced on a PacBio Sequel IIe platform at Berry Genomics (Beijing, China) following manufacturer’s instructions. Etiolated 2-week-old seedlings were collected and used for crosslinking, proximity ligation, and library construction. The Hi-C library prepared by Biozeron (Shanghai, China) and sequenced at the Illumina NovaSeq platform with paired-end 150 bp sequencing mode.

### Genome size estimation

To estimate the genome size of chia, 21 bp k-mer frequency of the PacBio HiFi reads was firstly counted with jellyfish (version 2.3.0) (Marcais and Kingsford, 2011). The k-mer frequency table was then used as input for GenomeScope2 (version 2.0) (Ranallo-Benavidez et al., 2020) to fit a diploid mathematical model to estimate the genome size, heterozygosity, and repetitiveness **(Supplemental Figure 3)**.

### Genome assembly

To assemble the nuclear genome using HiFi reads, three state-of-the-art genome assemblers were tested, including Flye (version 2.9) (Kolmogorov et al., 2019), HiCanu (version 2.2) (Nurk et al., 2020), and hifiasm (version 0.16.1) (Cheng et al., 2021). Flye applied a data structure of repeat graph (Kolmogorov et al., 2019). HiCanu was a modification of the Canu assembler (Koren et al., 2017) that designed for HiFi reads with homopolymer compression, overlap-based error correction, and aggressive false overlap filtering (Nurk et al., 2020). Hifiasm is a genome assembler specifically designed for HiFi reads (Cheng et al., 2021). The previous estimated genome size by GenomeScope2 was used as input parameter for Flye and HiCanu. While hifiasm do not require pre-estimated genome size. The results indicated that hifiasm with default parameters performed the best in terms of contiguity **(Supplemental Table 6)** and accuracy (**Supplemental Figure 16**).

To assemble the chia plastid genome, the GetOrganelle software (version 1.6.2) (Jin et al., 2020), which performs well in a systematic comparison of chloroplast genome assembly tools, (Freudenthal et al., 2020) was used. GetOrganelle firstly extracted Illumina short reads that could be mapped to the embryophyte plastomes (a library composed of 101 plastid genomes) by bowtie2 (version 2.3.4.1) (Langmead and Salzberg, 2012) and then assembled them using SPAdes (version 3.13.0) (Bankevich et al., 2012). GetOrganelle produced three contigs representing the large single copy (LSC), small single copy (SSC) and inverted region (IR) of the chia plastid genome. Such three contigs were then aligned against the plastid genome of *Salvia miltiorrhiza* (accession number: NC_020431.1) (Qian et al., 2013), a close relative of chia. The alignment was performed with minimap2 (version 2.11) (Li, 2018) and visualized with D-Genies (version 1.3.1) (Cabanettes and Klopp, 2018). The three contigs were then ordered into a complete plastid genome using a customized Perl (version 5.34.0) script based on the BioPerl toolkit (version 1.7.4) (Stajich et al., 2002). Next, CHLOË (version 7c33699, https://chloe.plastid.org/) was used for the annotation of protein-coding genes, transfer RNAs, and ribosomal RNAs in the plastid genome.

To obtain the chia mitochondrial genome, we inspected contigs produced by hifiasm and found contig ptg000033c (length: 313,444 bp, read depth: 557) was circular and had the highest average read depth. Then we submitted this contig to the AGORA web tool (Jung et al., 2018) for genome annotation with the protein-coding and rRNA genes of the *Salvia miltiorrhiza* mitochondrial genome (accession number: NC_023209.1) as reference. The results of AGORA were then manually corrected by 1) removing protein-coding genes less than 30 amino acids, 2) removing protein-coding genes with pre-stop codons, 3) correcting mislabeled positions of ribosomal RNA genes. The chia mitochondrial genome was then visualized using OrganellarGenomeDRAW (OGDraw, version 1.3.1) (Greiner et al., 2019).

The “1-to-1” coverage and identity rate of contigs against the chia plastid and mitochondrial genomes were calculated using the dnadiff program of the MUMmer package (version 3.23) (Kurtz et al., 2004).

To obtain chia pseudochromosome sequences, the top 21 contigs in length and the Hi-C data was used for scaffolding. Illumina sequencing adapters and low-quality sequences of Hi-C data were trimmed by trim_galore (version 0.6.7, https://github.com/FelixKrueger/TrimGalore) with default parameters (quality score: 20; minimum length: 20 bp), which is a wrapper of cutadapt (version 3.4) (Martin, 2011). The clean Hi-C data were analyzed by Juicer (version 1.6) (Durand et al., 2016), which produced high-quality DNA contact information. Then the 3D-DNA pipeline (version 180922) (Dudchenko et al., 2017) was used for ordering the contigs into pseudochromosomes. After visualizing the Hi-C contact map by Juicebox (version 1.9.1) (Durand et al., 2016), we manually connect the contigs using “run-asm-pipeline-post-review.sh” of the 3D-DNA pipeline to avoid splitting the contigs.

### Identification of rRNA repeats and telomere signatures

To predict the location of ribosomal RNA (rRNA) in the nuclear genome, Basic Rapid Ribosomal RNA Predictor (barrnap, version 0.9, https://github.com/tseemann/barrnap) was used, which using the nhmmer (version 3.1b1) (Wheeler and Eddy, 2013) to search the potential location of eukaryotes rRNA genes (5S, 5.8S, 28S, and 18S).

The telomere signature was examined using the program FindTelomeres (https://github.com/JanaSperschneider/FindTelomeres), which was a Python script for finding telomeric repeats (TTTAGGG/CCCTAAA). The results were further confirmed by TRF (version 4.09.1) (Benson, 1999) with parameters of “2 7 7 80 10 50 500 -m -d -h”.

Genome circular plots were created in Circos (version 0.69.6) (Krzywinski et al., 2009).

### Genome quality evaluation

The quality of the genome assembly was evaluated using three methods, including Benchmarking Universal Single-Copy Orthologs (BUSCO) (version 5.0.0) (Manni et al., 2021), LTR Assembly Index (LAI) (version 2.9.0) (Ou et al., 2018) and Merqury (version 1.3) (Rhie et al., 2020). Merqury is a tool for reference-free assembly evaluation. Additionally, Illumina short reads were mapped to chia genome assembly using bwa-mem (version 0.7.17) (Li, 2013). The mapping rate and error rate of the Illumina short reads were estimated by SAMtools (version 1.15.1) (Li et al., 2009).

### Genome annotation

A combined method was used for chia gene prediction, including *ab initio* prediction, EST discovery and protein homology search. To predict gene models, we firstly masked the repeats using RepeatMasker (version 3.1.2-p1) (Tarailo-Graovac and Chen, 2009). A species-specific repeat library was constructed for RepeatMasker using Repeatmodeler2 (version 2.0.2) (Flynn et al., 2020) and LTR_retriever (version 2.9.0) (Ou and Jiang, 2018). The LTR candidates for LTR_retriever was identified by LTR_FINDER_parallel (version 1.1) (Ou and Jiang, 2019) and LTRharvest (version 1.6.0) (Ellinghaus et al., 2008).

LTR_FINDER_parallel is a parallel wrapper of LTR_FINDER (version 1.07) (Xu and Wang, 2007). The chia transcriptome of 13 tissue types (involved seeds, cotyledon, shoots, leaves, internodes, racemes, and flowers) (Gupta et al., 2021) were retrieved from the NCBI SRA database (accession number: PRJEB19614) and *de novo* assembled using Trinity (version 2.11.0) (Grabherr et al., 2011). The assembled transcripts were used as expressed sequence tags (EST) evidence for further gene model prediction. Seven sets of protein sequences downloaded from public databases were used as protein homology evidences, including *Arabidopsis thaliana* (version Araport11) (Cheng et al., 2017), *Antirrhinum majus* (version IGDBV1) (Li et al., 2019), *Callicarpa americana* (Hamilton et al., 2020), *Salvia miltiorrhiza* (version 1.0) (Song et al., 2020), *Salvia splendens* (Dong et al., 2018), *Tectona grandis* (Zhao et al., 2019) and the UniprotKB/Swiss-Prot dataset (version release-2020_04) (Poux et al., 2017).

Maker (version 3.01.03) (Campbell et al., 2014) was run three rounds to train AUGUSTUS (version 3.4.0) (Stanke and Waack, 2003) and SNAP (version 2006-07-28) (Korf, 2004) gene prediction parameters. GeMoMa (version 1.8) (Keilwagen et al., 2019) and MetaEuk (release 5) (Levy Karin et al., 2020) were used with the above mentioned protein homology datasets to discover gene models. Finally, EVidenceModeler (EVM, version 1.1.1) (Haas et al., 2008) was used to combine all the above gene prediction evidences. The est2geome and protein2genome features produced by Maker were used as transcript and protein evidence for EVM. The AUGUSTUS and SNAP gene models were used as *ab initio* prediction evidence for EVM. The GeMoMa and EetaEuk produced gene models were used as OTHER_PREDICTION evidence, which means they do not provide an indication of intergenic regions (Haas et al., 2008). Gene function annotation was performed by InterProScan (version 5.52-86.0) (Jones et al., 2014) and AHRD (version 3.3.3) (Boecker, 2021).

### Genome evolution

Orthofinder (version 2.5.4) (Emms and Kelly, 2019) was used for the construction of orthologous groups. The STAG algorithm (Emms and Kelly, 2018) implemented in Orthofinder was used to estimate the species tree. Chia and other nine genomes were used for the construction of orthologous groups, including *Arabidopsis thaliana* (version Araport11) (Cheng et al., 2017), *Solanum lycopersicum* (version ITAG4.0) (Hosmani et al., 2019), *Antirrhinum majus* (version IGDBV1) (Li et al., 2019), *Tectona grandis* (Zhao et al., 2019), *Callicarpa americana* (Hamilton et al., 2020), *Jacaranda mimosifolia* (Wang et al., 2021), *Salvia bowleyana* (Zheng et al., 2021), *Salvia miltiorrhiza* (version 1.0) (Song et al., 2020), and *Salvia splendens* (version SspV2) (Jia et al., 2021). Gene family size expansion and contraction analysis was performed by CAFE5 (version 5.0.0) (Mendes et al., 2020). Synteny analysis was performed by the Python version of MCScan (version 1.1.17) (Tang et al., 2008). ParaAT (version 2.0) (Zhang et al., 2012) was used for prepare the alignment data for calculating Ks values, which was a wrapper of MUSCLE (version 3.8.1551) (Edgar, 2004) and PAL2NAL (version 13) (Suyama et al., 2006). KaKs_Calculator (version 2.0) (Wang et al., 2010) was used for calculating the Ks values using the YN model (Yang and Nielsen, 2000).

### Gene expression analysis

Besides the chia transcriptome of 13 tissue types that retrieved from the NCBI SRA database (accession number: PRJEB19614) (Gupta et al., 2021), another set of transcriptome data for chia seed development was retrieved from the NCBI SRA database (accession number: PRJNA196477), which was sampled in 3, 7, 14, 21, and 28 DAF (Sreedhar et al., 2015). The raw RNA-seq data that downloaded from the NCBI SRA database were firstly converted to FASTQ format using the fastq-dump command from the SRA Toolkit package (version 2.9.3, https://github.com/ncbi/sra-tools). The data were then trimmed using trim_galore and then mapped to the chia reference genome by STAR (version 2.7.5c) (Dobin et al., 2013). Gene counts were summarized by featureCounts (version 2.0.1) (Liao et al., 2014). FPKM values were calculated using functions of the DESeq2 package (version 1.32.0) (Love et al., 2014) in the R platform (version 4.1.1) (R Core Team, 2021).

### Multiple sequence alignment and phylogenetic tree construction

Visualization of multiple sequence alignment of the *FAD2, FAD3, FAD7, and FAD8* genes was performed using the Clustal Omega web tool (https://www.ebi.ac.uk/Tools/msa/clustalo/). Phylogenetic trees of the *FAB2*/*AAD*, *FAD2, FAD3*, *FAD7* and *FAD8* were constructed with the maximum likelihood method by IQ-TREE2 (Minh et al., 2020). The best-fitting amino acid substitution model was determined by ModelFinder (Kalyaanamoorthy et al., 2017).

### Data availability

The genome assembly and corresponding sequencing data were deposited at NCBI under BioProject number PRJNA864090 and at NGDC under accession number PRJCA010915.

## Author contributions

L.L. performed data analyses; J.S., M.Z., and S.I. prepared plant materials; S.I., Y.L., He.Z., Hu.Z. designed the project; L.L. and He.Z. wrote the manuscript.

## Acknowledgements

We thank the Core Facility for Genomics of Shanghai Center for Plant Stress Biology (PSC) for the construction of the Illumina sequencing library and the Core Facility for Bioinformatics of PSC for the maintenance of the high-performance computing (HPC) clusters. The authors declare no conflict of interest.

